# Auditory evoked potentials to changes in speech sound duration in anesthetized mice

**DOI:** 10.1101/329433

**Authors:** A. Lipponen, J.L.O. Kurkela, Kyläheiko I., Hölttä S., T. Ruusuvirta, P. Astikainen

## Abstract

Electrophysiological response termed mismatch negativity (MMN) indexes auditory change detection in humans. An analogous response, called the mismatch response (MMR), is also elicited in animals. Mismatch response has been widely utilized in investigations of change detection in human speech sounds in rats and guinea pigs, but not in mice. Since e.g. transgenic mouse models provide important advantages for further studies, we studied processing of speech sounds in anesthetized mice. Auditory evoked potentials were recorded from the dura above the auditory cortex to changes in duration of a human speech sound /a/. In oddball stimulus condition, the MMR was elicited at 53-259 ms latency in response to the changes. The MMR was found to the large (from 200 ms to 110 ms) but not to smaller (from 200 ms to 120-180 ms) changes in duration. The results suggest that mice can represent human speech sounds in order to detect changes in their duration. The findings can be utilized in future investigations applying mouse models for speech perception.

## 1. Introduction

Human speech sounds are spectrotemporally complex. They must be successfully processed by the brain to encode phonemes, syllables, words or sentences. Temporal features of the speech sounds are especially important, since speech recognition is preserved in humans even when spectral information is impoverished as long as temporal cues remain available (Shannon, Zeng, Kamath, Wygonski, & Ekelid, 1995). Moreover, vowel durations play a critical role in determining semantics in quantitative languages.

Vowel duration processing can be addressed via an electrophysiological response called mismatch negativity (MMN) (Näätänen, Gaillard, & Mäntysalo, 1978); for a review see Pulvermüller & Shtyrov, 2006). In humans, the MMN is observed in scalp-recorded event-related potentials at 100-250 ms after the rare ‘deviant’ stimulus that occasionally replaces the occurrences of the frequently presented‘standard’ stimulus (Näätänen et al., 1978). Importantly, the MMN is elicited only when deviant sound input is detected to violate the regularity formed by the standard stimuli (Näätänen, Jacobsen, & Winkler, 2005), not for instance by the first stimulus in a sound series (Näätänen et al., 2005).

Mismatch response (MMR) analogous to the MMN has been found in several animal species (e.g. in cat Csépe, Karmos, & Molnár, 1987; in monkey Javitt, Schroeder, Steinschneider, Arezzo, & Vaughan, 1992; in guinea pig Kraus et al., 1994; in mice Umbricht, Vyssotki, Latanov, Nitsch, & Lipp, 2005), but most of the animal studies are conducted in rats (e.g. Ruusuvirta, Penttonen, & Korhonen, 1998; Eriksson & Villa, 2005; von der Behrens, Bäuerle,Kössl, & Gaese, 2009; Tikhonravov et al., 2010; Astikainen et al., 2011; Astikainen, Ruusuvirta, Wikgren, & Penttonen, 2006; Nakamura et al., 2011; Klein, von der Behrens, & Gaese, 2014, for a review see Grimm, Escera, & Nelken, 2016). Similarly to the human MMN (Näätänen et al., 2005), most of the studies support the view that MMR in rats reflects detection of regularity violations (Ahmed et al., 2011; Astikainen et al., 2011; Jung et al., 2013; Nakamura et al., 2011; Parras et al., 2017; Shiramatsu, Kanzaki, & Takahashi, 2013); for a review see (Harms, Michie, & Näätänen, 2016). Namely, MMR disappears in a stimulus condition where the regular standard sound is replaced with a set of constantly varying sounds (so-called equiprobable condition, also known as many standards condition, (Jacobsen & Schröger, 2001; Schröger & Wolff, 1996)). However, some studies have suggested that also neural adaptation, which is stronger for the responses to repetitive standard sounds than rare deviants, could explain the differential response for the deviant sounds, although in these studies the control conditions were different from equiprobable condition (Eriksson & Villa, 2005; Lazar & Metherate, 2003). There is only one MMR study conducted in mice where the equiprobable control condition has been applied (Kurkela, Lipponen, Kyläheiko, & Astikainen, 2018). It demonstrated a MMR to frequency changes which reflected regularity violations starting at 53.5 ms latency, but not in the earlier latency.

Recordings of the MMR have been used to index processing of human speech sounds in anesthetized guinea pigs and rats (Kraus et al., 1994). These studies have investigated change detection in syllables (Ahmed et al., 2011; Kraus et al., 1994; McGee et al., 2001) and syllable patterns (Astikainen, Mällo, Ruusuvirta, & Näätänen, 2014). However, no MMR studies applying spectrotemporally complex, such as human speech sounds have been conducted in mice. A mice model for complex sounds human speech perception would be particularly valuable since there are transgenic mice strains available. Here we investigated detection of changes in vowel duration in anaesthetized mice. Auditory evoked potentials were recorded from the dura above the auditory cortex. Duration changes were presented in 8 separate oddball conditions in which a deviant sound either of 110, 120, 130, 140, 150, 160, 170 or 180 ms in duration was interspersed with a 200-ms standard sound. Only decrements in the speech sound duration were applied, because increments in duration would suffice to elicit larger responses due to higher stimulus energy delivered by deviants than standards. In addition to the oddball conditions, an equiprobable control condition was applied to probe into whether MMR, if observed, reflects mechanism related to detection of regularity violations or whether stimulus-rate related explanations (e.g., release for neural refractoriness, (Näätänen et al., 2005) would suffice for accounting for response elicitation.

## 2. Material and methods

### 2.1 Animals

The experiments were approved by the Finnish National Animal Experiment Board (ESAVI/10646/04.10.07/2014). They were carried out in accordance with the European Communities Council Directive (86/609/EEC) on the care and use of animals in experimental procedures. Eight male (n = 8) C57Bl/6J mice from the Animal Center of UEF, Kuopio were used in the experiment (weight 24.8 ± 2.0 g, age 12.1 ± 1.4 weeks, mean ± SEM). The animals were group housed with water and fed ad libitum. The mice were in a controlled environment (constant temperature 22 ± 1°C, humidity 50–60%, lights on 07:00–19:00 h). At the end of the experiment anesthetized animals were sacrificed with cervical dislocation.

### 2.2 Surgery and LFP recording

For the surgery and acute recordings, the animals were anaesthetized at first with isoflurane (initial dose 5 % in 1 l/min of pressure air followed with maintaining dose of 1-2 % in 1 l/min of pressure air). As soon as the animal was head fixed to the stereotactic device (Kopf, CA, USA) a single dose of urethane (Sigma-Aldrich, St. Louis, MO, USA; 7,5 g/100 ml-solution) was administered i.p. (1.2 g/kg) and administration of isoflurane was gradually diminished over 5 min (Barth & Mody, 2011). The level of anesthesia was controlled by regular testing of pedal withdrawal reflex and if required, extra doses (0.1–0.2 ml) of urethane were given.

Under full anesthesia skin and muscle tissue over the skull were removed. For the reference electrode, a hole was drilled in the skull over the right side of the cerebellum and a small insulin needle (BD Lo-Dose syringe, USA) was inserted in the cerebellum as reference electrode (AP −5.8 mm, ML: 2–3 mm and DV: 2 mm). A similar needle electrode inserted subcutaneously into the neck served as the ground electrode. Next, the upper half of the squamosal skull bone was removed over revealing the left primary auditory cortex but the dura was left intact (Stiebler, Neulist, Fichtel, & Ehret, 1997).

A tip of a Teflon-insulated silver wire (diameter 200 µm, A-M Systems, Carlsberg, WA, USA) was placed on the surface of the dura above the auditory cortex (2.2–3.2 mm posterior and 4–4.5 mm ventral to bregma). Although the coordinates for the craniotomy refer to the primary auditory cortex, some activity from the higher sensory areas may also have been captured. The latter is due to individual differences in the organization of the primary auditory cortex and the relatively large size of the tip of the electrode, allowing the signal to be conducted from adjacent areas. To reduce the variability the electrode location was also guided by on-line recorded auditory evoked potentials in response to 200-ms stimuli (presented as the standard tone in the actual experiment).

A continuous electrocorticogram was first 10-fold amplified using a low-noise MPA8I pre-amplifier (MultiChannel Systems MCS GmbH, Reutlingen, Germany). The signal was further fed to a filter amplifier (FA64I, filter: 1–5000 Hz, MCS). All the signals were digitized (USBME-64 System, MCS) and recorded with McRack software (MCS), using a 2000-Hz sampling rate. Finally, all the signals were digitally band-pass filtered between 1 and 500 Hz (high-pass: low-pass: fourth-order Bessel).

### 2.3 Stimuli

The speech sound /a/ used in the experiment was originally recorded at sampling rate of 44.1 kHz from a female native Finnish speaker. The sound was then digitally edited using SoundForge software (SoundForge 10, Sony Corporation, Japan) to ensure it had a constant duration of 200 ms. The 200-ms speech sound was digitally shortened in 10-ms steps and a 5-ms fade-out was added in each sound in Soundforge Pro 10.0 (MAGIX Software GmbH, Germany).

This resulted in the 10 speech sounds used in the experiment (200, 190, 180, 170, 160, 150, 140,130, 120 and 110 ms in duration, figure 1).

**Figure 1.**
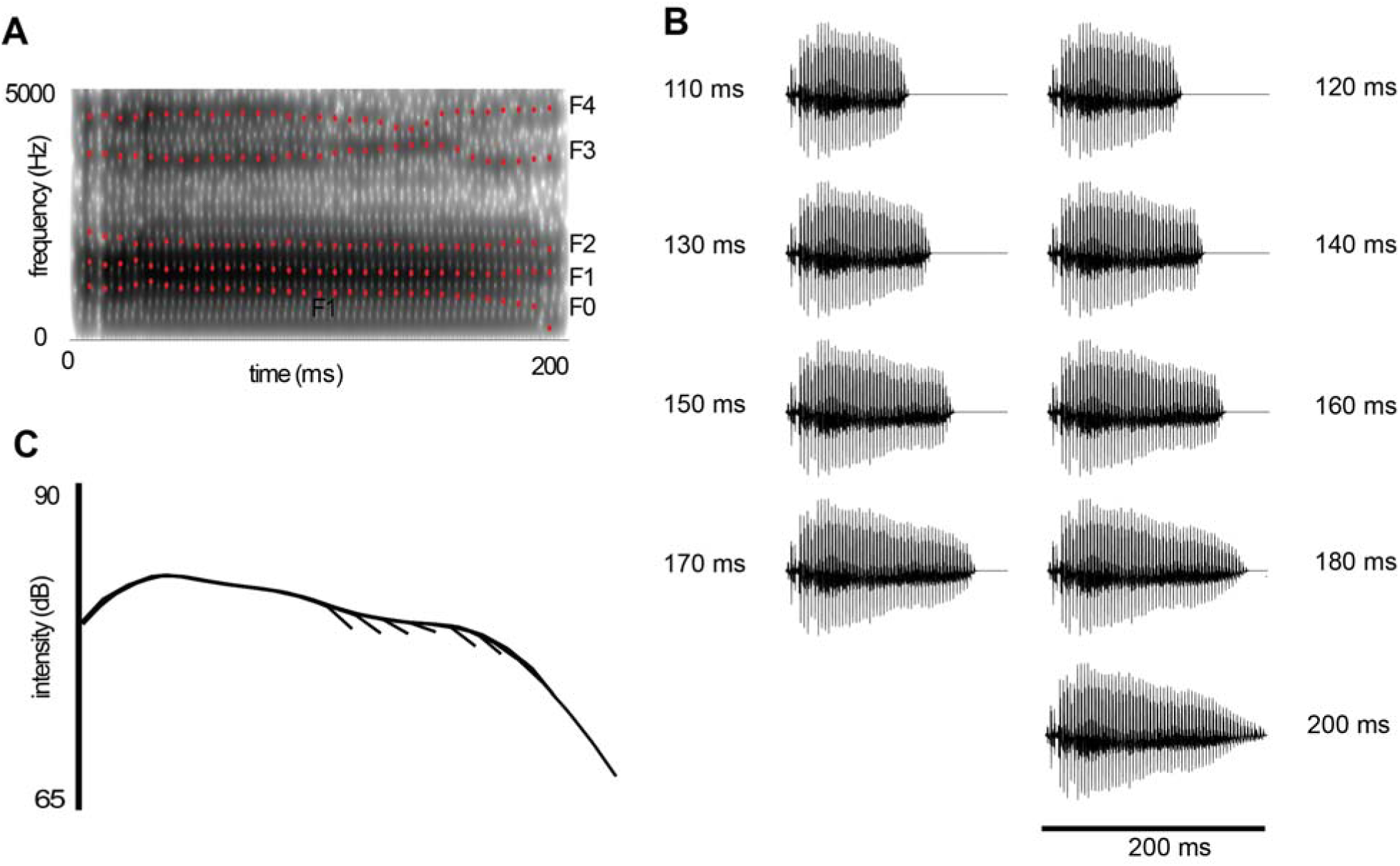
Human speech sounds used in the experiment. (A) The spectrogram and F0, F1, F2, F3 and F4 formants of the standard speech sound /a/. (B) The time-amplitude patterns of the sounds used in the experiment (110, 120, 130, 140, 150, 160, 170, 180 and 200 ms), and (C) the intensity of the same sounds. The intensity of all the sounds is the same except the last 5-ms fade-out in each sound.

Sounds were presented via an active loudspeaker system (Studiopro 3, N-audio, Irwindale, CA, USA), and the stimulus presentation was controlled by E-prime 2.0 software (Psychology Software Tools, Pittsburg, PA). The stimulation was presented with the passive part of the loudspeaker system directed towards the right ear of the animal at a distance of 20 cm with sound pressure level (SPL) of 70 dB, as measured with sound level meter (c weighting, Sound level meter Type 2240 Brüel & Kjær Sound & Vibration, DK-2850, Nærum, Denmark).

Two different types of stimulus conditions were presented to the subjects: the equiprobable condition and the oddball condition (figure 2). In one stimulus block the equiprobable condition was applied. In it, all the 10 speech sounds were presented with a same probability (p = 0.10) in random order and with the inter-stimulus interval (ISI) of 400 ms. Nine stimulus blocks of oddball condition were applied in which the deviant duration was 110, 120, 130, 140, 150, 160, 170, 180 and 190 ms, but due to a technical failure, the oddball stimulus block with the 190-ms deviant tone is not reported. In each oddball condition, the 200-ms speech sound served as a repeatedly presented standard sound (p = 0.90) and one of the nine shorter sounds served as a deviant sound (p = 0.10). The deviant sound was delivered among the standard sounds in random order, but there were always at least two standard sounds between two deviant sounds. There were 1000 stimuli in each stimulus block and they were presented in a counterbalanced order between the subjects.

**Figure 2.**
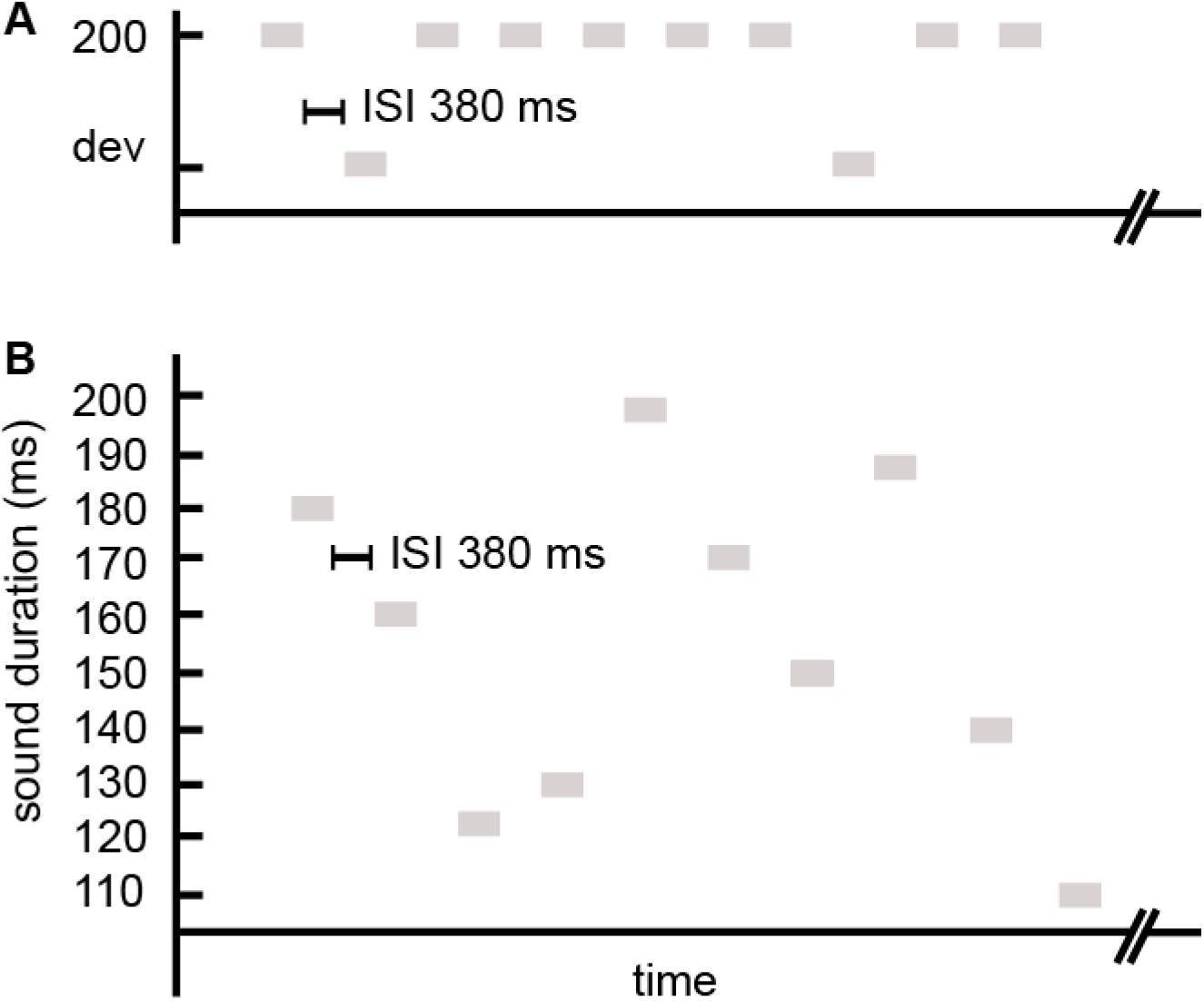
Stimulus series used in the experiment. The tones were presented in the oddball condition (A) and equiprobable condition (B). A) In the oddball condition, a deviant sound (*P*= 0.1) was interspersed with a repetitive standard sound (*P* = 0.9). B) In the equiprobable condition, control stimuli of 9 different frequencies were presented with the same probability (*p* = 0.1).

### 2.4 Data analysis

The data analyses were performed offline using Brain Vision Analyzer (Brain Products, Gilching, Germany), GraphPad Prism 5.03 (GraphPad Software) and Excel 2016 (Microsoft Office).

The data were segmented (from −50 to 400 ms from stimulus onset) separately for the responses to the deviant and standard sound immediately preceding the deviant sound in the oddball condition, and for responses to each tone in the equiprobable condition. Data segments were baseline corrected against the mean of the signal during a 50-ms time window prior to sound onset. Data segments were averaged for each animal separately for different deviant and standard tones preceding the deviant tones, and for each tone presented in equiprobable condition.

### 2.5 Statistical analysis

Paired t-tests were applied to compare the averaged response amplitudes in a point-by-point manner. The comparisons were first made between responses to standard and deviant sounds for each oddball condition. If the point-by-point comparison values reached statistical significance (p < .05) for at least 20 consecutive sample points, i.e. over a 10-ms time span, differential responses to oddball deviant sounds were considered to exist and not be accounted for by random signal fluctuations. In this case, the comparison was also made between oddball deviant response and control-deviant response (i.e. the same sounds presented in the equiprobable condition). Due to technical problems, recording of the equiprobable control condition in one animal was lost, and therefore the comparison of the responses in the oddball and control condition consists of seven animals. The t-tests were two-tailed when standard and deviant responses were compared in the oddball conditions, because we were not able to predict the direction of a possible differential response (since for example animal species, location of the reference electrode and anesthesia could affect the polarity of the differential response (Harms et al., 2016). The t-tests comparing deviant and control-deviant responses were one-tailed because we specifically tested, whether the responses were larger for the deviant than for the control deviant sounds (towards the polarity the deviant sounds displaced the response in comparison to the standard sound). Cohen’s d is reported for the effect size.

## 3. Results

Overall, the sounds evoked a prominent response of positive polarity peaking at ~35 ms after stimulus onset (Figure 2). Visual inspection revealed a positive displacement to the oddball deviant sounds in comparison to the 200-ms standard sounds, especially to the most distinct 110-ms deviant sound.

Point-by-point t-tests comparing the amplitude of the standard and the deviant sound responses revealed a statistically significant amplitude difference at 163-369 ms after stimulus onset (all *p* < 0.05) for the 110-ms deviant sound (corresponding to the latency of 53-259 ms after the change onset). The effect size was the largest at 227 ms latency (i.e. 117 ms after the change onset, d = 1.56). However, the other deviant sounds elicited no differential responses compared to the standard sound.

Next, the responses to the 110-ms deviant sound in the oddball condition and the responses to the corresponding control-deviant sound were compared within the time interval in which the differential (MMR) response to the deviant responses in the oddball condition. A marginally significant difference (*p* < 0.1) between the oddball sound and the control sound was found at 235-369 ms latency (corresponding to 125-259 ms post-change), the effect size being largest at the latency of 337 ms (i.e. 227 ms post-change, *P* = 0.06, d = 0.8) (figure 3).

**Figure 3.**
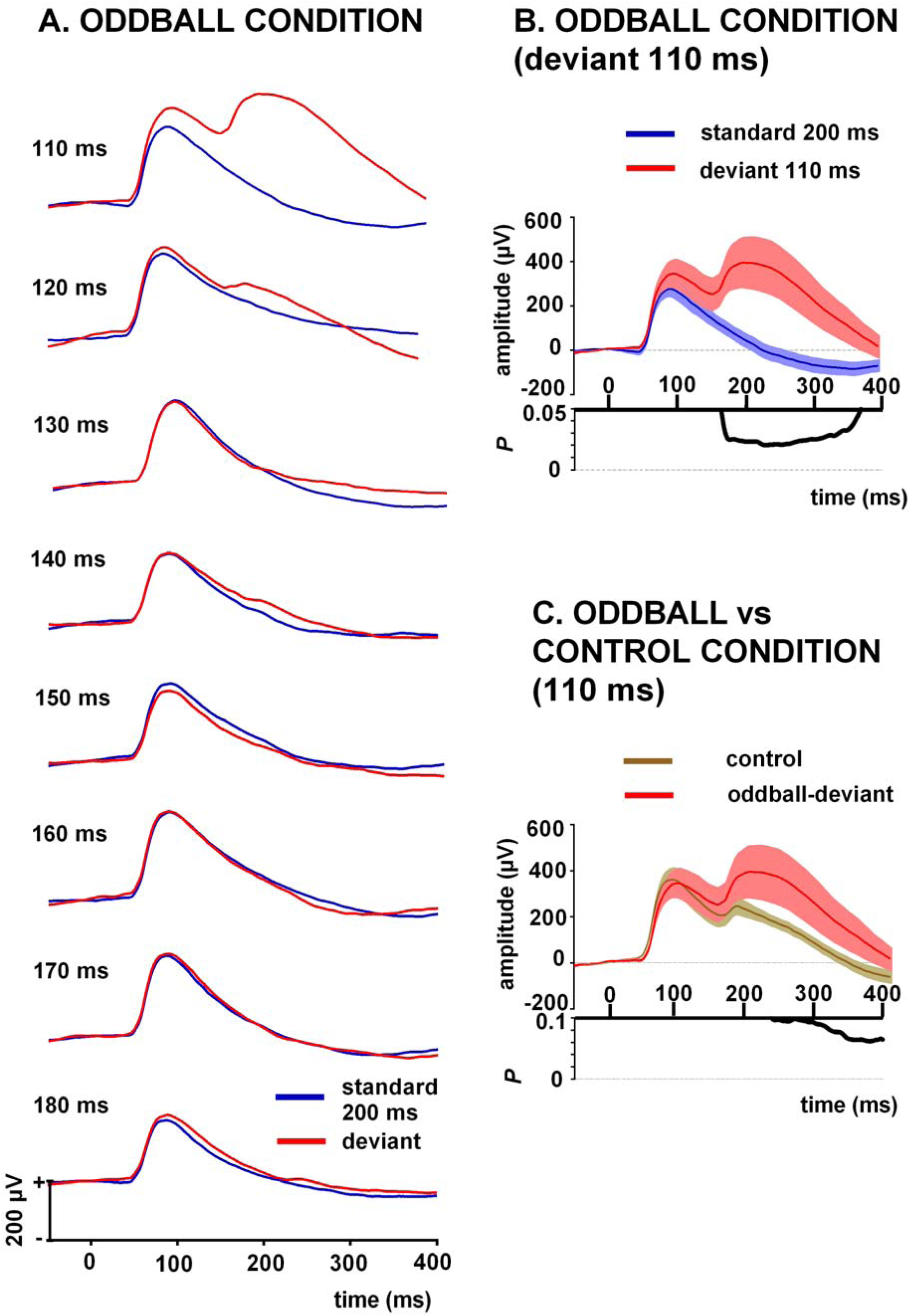
Grand-averaged responses recorded from the dura above the auditory cortex. A) Evoked potentials for the deviant (110, 120, 130, 140,150,160, 170 and 180 ms) and to the 200-ms standard sound. The sounds were presented in the oddball condition. B) Evoked potentials to the 110-ms deviant and 200-ms standard stimulus in the oddball condition. The point-by-point t-tests to response amplitudes revealed a significant difference between these responses at latency of 163-369 ms (corresponding to 53-259 ms after the change onset). C) Evoked potentials to the 110-ms deviant sound (oddball condition) and corresponding control sound (equiprobable condition). A marginally significant difference between these responses was found at 235-369 ms latency (p < 0.1; corresponding to 125-259 ms post-change). Stimulus onset at 0 ms. The waveforms are presented as mean (A) and mean ± SEM (B and C).

## 4. Discussion

We investigated whether the MMR, a non-human correlate of the human MMN indexing auditory change detection, is elicited in anaesthetized mice in response to duration changes in a human speech sound /a/. We therefore studied not only whether such changes elicit MMR-like waveform, but also whether this waveform reflects these changes as such rather than the lower presentation rate of the deviant sound relative to the standard sound (i.e. lower level of refractoriness or neural adaptation in deviant-related than standard-related neural pathways). This was allowed by the use of the equiprobable condition (Harms et al., 2016) as a control condition for the oddball condition.

We found that only the shortest (110-ms) deviant sound interspersed with the 200-ms standard sound (45 % duration decrease) elicited an observable MMR. Consistently, mice have been found to generate the MMR to 50 % shortenings (from 100 ms to 50 ms) of a repeated sinusoidal sound (Umbricht et al., 2005). Our result is also in line with a study in rats (Roger, Hasbroucq, Rabat, Vidal, & Burle, 2009) showing that a 33% change but not a 16 % change in the duration of a sinusoidal sound (100-ms deviant vs. 150-ms standard) elicited the MMR. It thus seems that, overall, duration changes could be considerably more difficult to detect by rodents than by humans who show robust MMNs to as small as a 10 % change in the duration of a sound (100 ms vs. 110 ms, Jaramillo, Paavilainen, & Näätänen, 2000). Note, however, that behaviorally measured duration discrimination in mice has been suggested to violate the Weber’s law (the perceived change in stimuli is proportional to the initial stimuli, Klink & Klump, 2004), so the findings of discrimination thresholds within one range of stimulus durations might not be applicable to stimulus durations outside this range.

We applied the equiprobable control condition to test whether the lower presentation rate of the deviant sound than the standard sound suffices for explaining the MMR in the oddball condition (May & Tiitinen, 2010). Responses to the 110-ms sound in the oddball condition were only marginally significantly (p < 0.1) higher in amplitude than those to the same sound in the control condition between 125 and 259 ms post-change. This set of findings suggests that the early part of MMR could be linked to stimulus presentation rate inducing lower levels of neural refractoriness or adaptation in deviant-than standard-related neural pathways rather than auditory change detection. the later part of the response showing more relevance to auditory change detection in particular. These findings corroborate previous observations of late (> 200 ms) rather than early MMRs to non-speech sounds as change-specific responses (Chen, Helmchen, & Lütcke, 2015), implying that speech sounds appear to be treated quite similarly to non-speech sounds in the mouse brain in auditory change detection tasks.

## Acknowledgments

We would like to thank Dr. Markku Penttonen for his valuable help in data collection phase and Professor Hua Shu’s research group in preparing the stimuli, and Mr. Petri Kinnunen for preparing the stimulus presentation. This work was supported by the Academy of Finland, grant number 273134 to Piia Astikainen. The funding source had no involvement in the study.

